# Methylartist: Tools for Visualising Modified Bases from Nanopore Sequence Data

**DOI:** 10.1101/2021.07.22.453313

**Authors:** Seth W. Cheetham, Michaela Kindlova, Adam D. Ewing

## Abstract

Methylartist is a consolidated suite of tools for processing, visualising, and analysing nanopore methylation data derived from modified basecalling methods. All detectable methylation types (e.g. 5mCpG, 5hmC, 6mA) are supported, enabling integrated study of base pairs when modified naturally or as part of an experimental protocol.

**Background:** Covalent modification of nucleobases is an important component of genomic regulatory regimes across all domains of life [1–3] and is harnessed by various genomic footprinting assays, including DamID[4], SMAC-seq[5], and NOMe-seq[6]. Nanopore sequencing offers comprehensive assessment of base modifications from arbitrarily long sequence reads through analysis of electrical current profiles, generally through machine learning models trained to discriminate between modified and unmodified bases [7]. An increasing number of computational tools have been developed or enhanced for calling modified bases [8], including nanopolish [7], megalodon [9], and guppy [10], along with an increasing number of available pre-trained models.

## Results and Discussion

Methylartist offers novel and useful visualisation outputs beyond those available through extant visualisation tools aimed at nanopore-derived methylation [11–13] in terms of the plots and options that it offers as well as support for arbitrary modification types. This has utility for identification of modified bases in assay-specific contexts which include GpC methylation (NOMe-seq), and 6mA (SMAC-seq, DamID in a 5′-GATC-3′ context, as well as native RNA base modifications). With few exceptions [14,15], most currently available models for calling modified bases involve some form of methylation or hydroxymethylation, so modifications will be referred to collectively as “methylation”, without loss of generality.

Modified bases are called from signal-level data stored in fast5 files using an appropriate basecalling model. Methylartist supports importing modified base calls from megalodon (db-megalodon), nanopolish (db-nanopolish) and guppy (db-guppy). Additional import methods for translating the basecalling output to the SQLite database format used by methylartist will be made available as the need arises. To demonstrate the capabilities of methylartist, we sequenced MCF-7 cells sourced from ATCC and from ECACC on the Oxford Nanopore Technologies PromethION platform. MCF-7 is a widely-studied breast cancer cell line with many sub-lines expressing different cellular phenotypes [16]. We anticipated that sourcing cells originating from different repositories would yield divergent yet comparable methylation profiles for demonstration purposes.

The command “methylartist segmeth” aggregates methylation calls over segments into a table of tab-separated values, useful for comparing whole-genome methylation or methylation over various annotations such as promoters, enhancers, or transposable element families. The resulting table is useful on its own or as input to “methylartist segplot” or “methylartist composite”. Category-based methylation data aggregated with “segmeth” can be plotted either as strip or violin plots using the “segplot” command (Figure 1).

**Figure 1:**
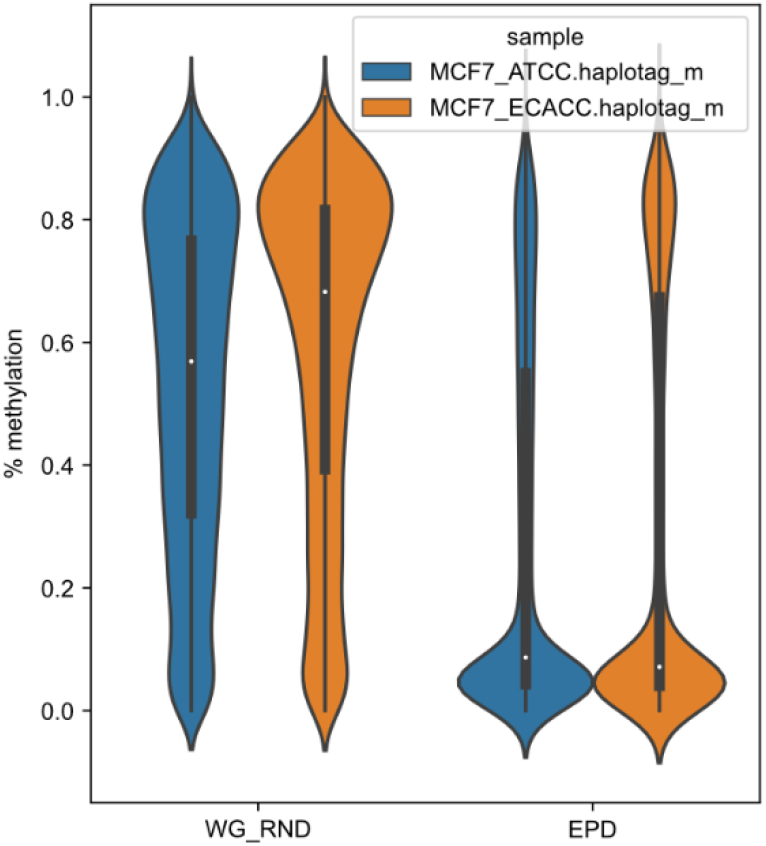
Violin plots from methylartist segplot showing the overall 5mCG distribution among 50000 random 590-bp segments of the genome (WG_RND) compared to a +/− 250bp window around eukaryotic promoter database (EPD) promoter annotations, which are 90bp.

Locus- or region-specific plots can be created in two ways, depending on the size of the window. For local regions, “methylartist locus” will generate plots similar to the example in Figure 2a, which depicts the methylation status of WNT7B known to be expressed in MCF-7 cells [17], and SHH, which appears to have a differentially methylated CpG island between the ATCC and ECACC cultivars (Figure 2b). These plots include an optional track showing genes, methylation calls relative to aligned read positions, a translation from genome space into a modified base space consisting only of instances of the methylated motif, a plot of the methylation statistic (e.g. log likelihood ratio), and a smoothed sliding-window plot showing methylation fraction across the region. The “locus” plotting mode also supports separating methylation profiles by phase, if the .bam files are first phased via WhatsHap [18] to add the “PS” and “HP” tags. Using this feature we can show apparent haplotype-specific methylation patterns that differ between the ATCC and ECACC cultivars in the TP53BP1 gene (Figure 2c).

**Figure 2:**
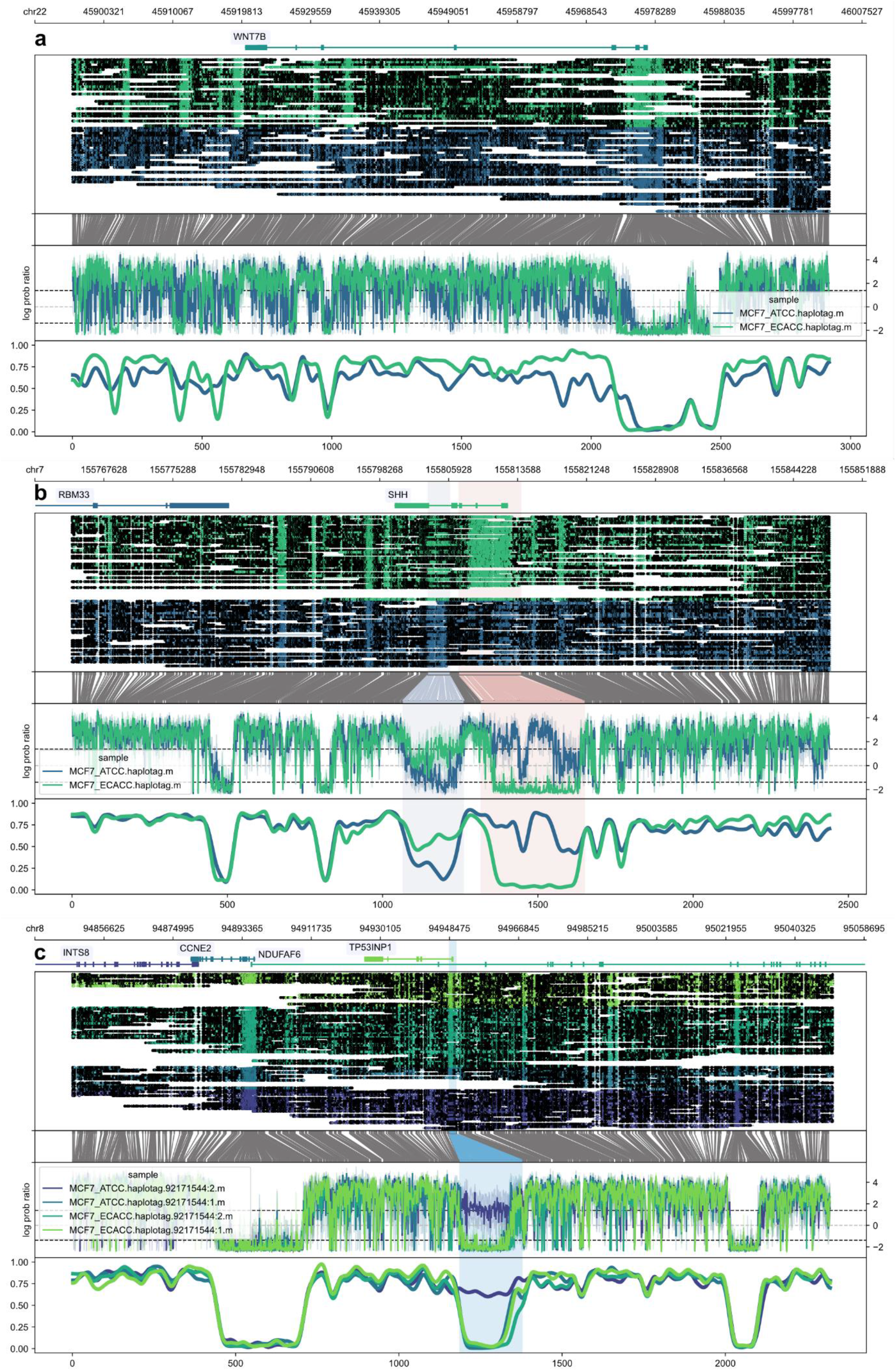
Methylartist locus plots showing methylation profiles for WNT7B **(A)**, SHH **(B)**, and allele-specific methylation profiles for TP53INP1 **(C)**. For each plot, the panels show the following information from top to bottom: genes (exons as boxes, introns as connecting lines) with optional labels, read alignments grouped and coloured by sample with methylation motifs (CpG) marked as open or closed dots, translation from genome coordinate space into a reduced modified base space (in these cases, CG dinucleotides), a “raw” plot of the methylated base statistic (in this case, log probability ratios), and finally a smoothed plot of the methylation profile. Plots **(B)** and **(C)** demonstrate the use of one or more highlights, which can be used to indicate regions of interest (in these cases, selected CpG islands).

For larger regions, “methylartist region” may yield a more expedient result as it aggregates methylation calls into bins, which can be normalised for occurrences of the methylation motif per-bin. The meaning of “larger” here depends on the density of methylation motifs in the region but generally 800kbp-1Mbp is a reasonable threshold. Region plots can also span an entire chromosome efficiently (Figure 3). The presentation is similar to “methylartist locus” but without the methylation statistic plot, and the coordinate transformation is from genome space into binned modified base space where bin sizes are normalised to equalise content of the modified base motif. Unless overridden via other options, the alignment plot is removed for regions larger than 5 Mbp and the panels rescaled appropriately. Both locus and region plots support an extensive set of parameters controlling dimensions, colour selection, highlighting, smoothing parameters, and panel ratios.

**Figure 3:**
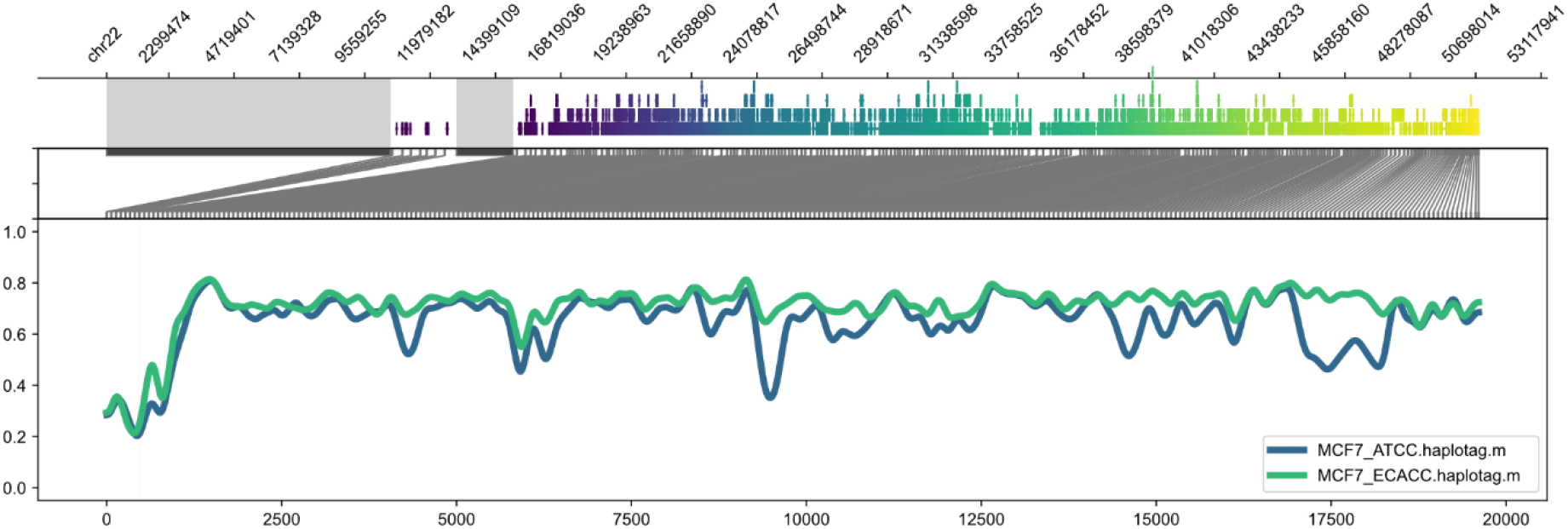
Demonstration of a larger scale methylartist region plot comprising all of human chromosome 22. The content of the plot is as described for Figure 1 but without the read alignment or raw statistic plots. The regions annotated as centromeric are shaded in grey (human chr22 is acrocentric). Coordinates across the bottom refer to methylation bins used in the smoothed methylation profile plot.

To demonstrate analysis and plotting of non-CpG methylation in a relevant context, we carried out a version of SMAC-seq [5], in which nuclei are treated with EcoGII [19] to mark accessible chromatin with 6mA. We analysed the SMAC-seq data with megalodon using the “res_dna_r941_min_modbases-all-context_v001” model from the rerio repository (https://github.com/nanoporetech/rerio), created a methylartist database via “db-megalodon” and identified loci from the eukaryotic promoter database (EPD) [20], with high 6mA relative to unmodified adenine using the “segmeth” utility in methylartist. In general, we see higher apparent 6mA methylation in regions defined by the Eukaryotic Promoter Database as compared to 50k size-matched regions of the genome drawn at random (Supplemental Figure 1). These loci were plotted en masse via the “locus” tool and screened visually, examples with corresponding CpG methylation plot are shown in Figure 4 and Supplemental Figure 2. Methylartist supports settings to improve visualisation of data where the expected distribution is a series of peaks, including the ability to limit inclusion of reads with an unusually high fraction of methylated bases (--maxfrac), and the option to skip sites below a threshold of methylated + unmethylated call coverage (--mincalls).

**Figure 4:**
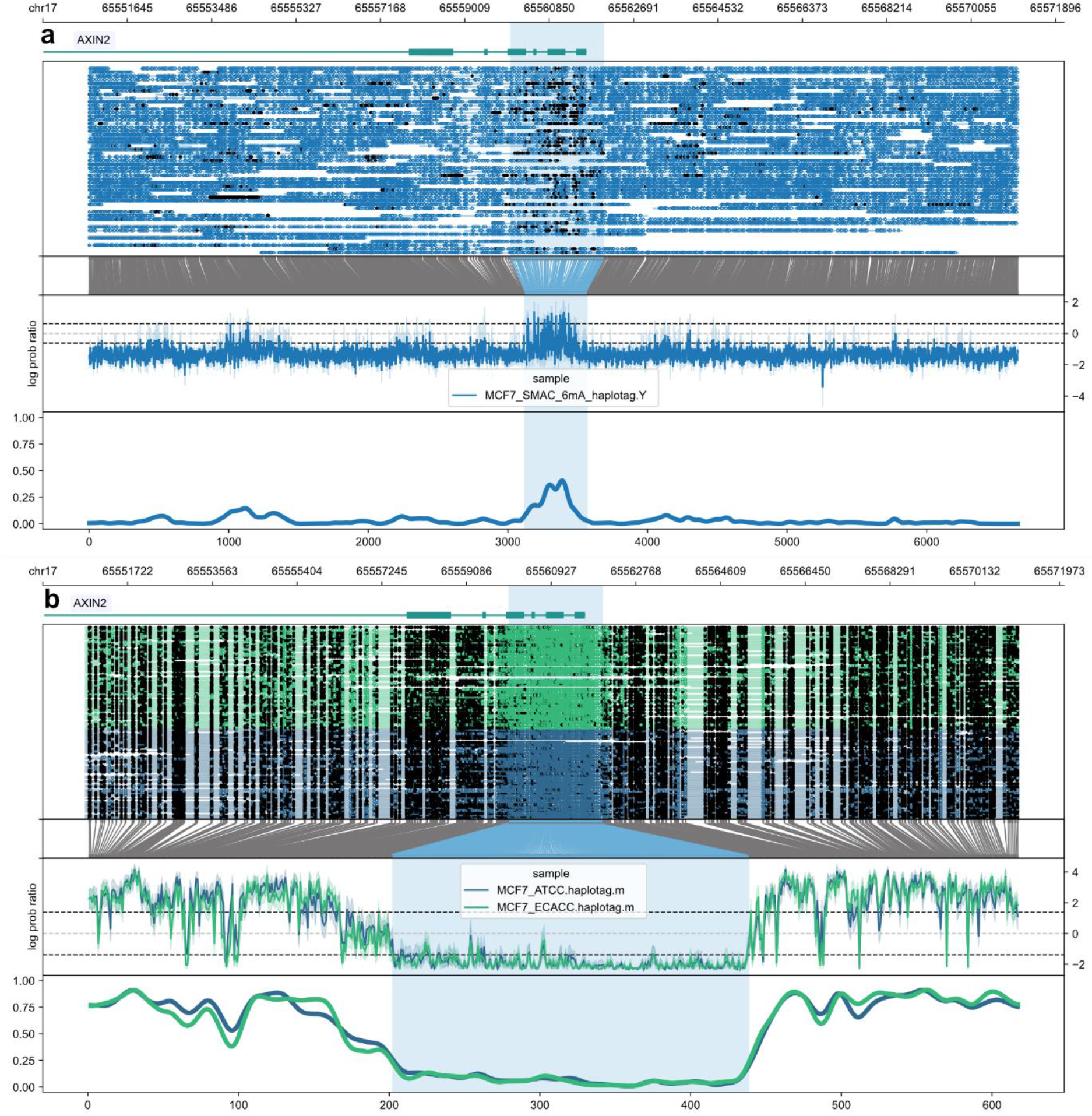
Example plot incorporating non-CpG methylation comparing the 6mA profile for an AXIN2 promoter from SMAC-seq on ATCC MCF-7 cells **(A)** with the 5mCG methylation profile from both MCF-7 cultivars used in this study **(B)**. Plots are as described in Figure 1, substituting A for CG in the modified base space in plot (A). The highlighted region, which is the same in both plots, corresponds to increased 6mA indicating accessible chromatin in plot **(A)** and decreased 5mCG consistent with regulatory elements in plot **(B)**.

In order to facilitate the study of methylation patterns across families of highly duplicated sequences such as transposable elements [21], methylartist supports a “composite” methylation plot, which aligns each instance of a repeat element family to a user-supplied consensus sequence and shows the methylation profile of a user-defined number of individual elements (Figure 5). Finally, the “wgmeth” tool in methylartist can also output bedMethyl files and files suitable for input to DSS, a package for assessing differential methylation [22].

**Figure 5:**
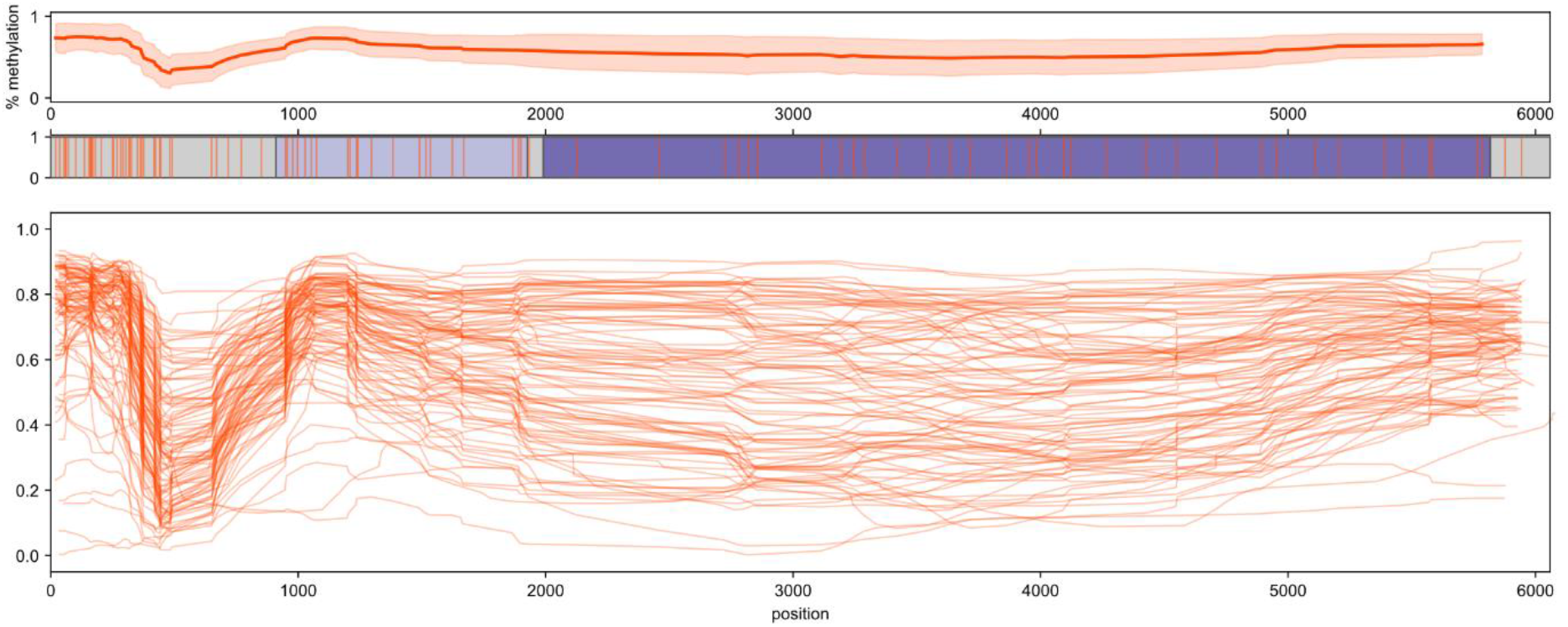
Repetitive element composite methylation profiles. Methylation (5mCG) profiles of full length (∼6000bp) human-specific LINE-1 elements in ECACC MCF-7 cells using the methylartist composite function. From top to bottom the elements of this plot include an indication of the average methylation level across the L1 consensus element, a plot indicating features (ORF1, ORF2) of the L1 consensus sequence (controlled through a table input to methylartist, orange vertical lines = CpGs), and a plot of the methylation profiles of 300 individual full-length L1 elements, randomly selected.

## Conclusion

We have demonstrated that methylartist has substantial utility as a plotting tool and as an accessible augmentation to the available tools for analysis and visualisation of nanopore-derived methylation data, including of non-CpG methylation useful in chromatin footprinting assays. Functionality will be expanded and updated in the future as use cases arise and as methods for analysis of nanopore data continue to evolve. For instance, the ability to seamlessly compare the same dataset on multiple modified basecallers or models could be useful for benchmarking applications. For demonstration purposes, we sequenced MCF-7 cultivars from two sources (ATCC, ECACC). While a comprehensive assessment of differences between cultivars is beyond the scope of this paper, it is well documented that significant differences exist between MCF-7 sub-lines [16], which is reflected in some of the examples used here. Methylartist provides a set of readily usable computational tools with which comprehensive assessment and visualisation of inter- and intra-sample methylation is possible including allele-specific methylation (Figure 2c), and methylation in regions difficult to comprehensively access with short-read or hybridisation-based methods such as transposable elements [21].

## Methods

### Cell culture

MCF-7 cells (ATCC, ECACC) were grown to 60-80% confluency in high-glucose Dulbecco’s Modified Eagle Medium (DMEM, Life Technologies) supplemented with 10% heat-inactivated Fetal Bovine Serum (Life Technologies), 2mM L-Glutamine (Life Technologies) and 100U/mL Penicillin-Streptomycin solution (Life Technologies). Cells were washed with Dulbecco’s Phosphate Buffered Saline (DPBS, Life Technologies), lifted with Trypsin 0.25% EDTA (Life Technologies), pelleted by centrifugation, and washed again with DPBS.

### Long-read PromethION sequencing

Genomic DNA was isolated using a Circulomics Big DNA Tissue Kit. Both the high molecular weight (HMW) and ultra-high molecular weight (UHMW) protocol was carried out for each cultivar (ATCC, ECACC) according to the manufacturer’s instructions for a total of 4 PromethION flow cells (Supplemental Table 1). Due to high DNA viscosity the UHMW protocol was modified to include vortexing after addition of CLE3 digestion buffer and incubation at 37°C instead of RT. For simplicity and for demonstration purposes replicates were combined across cultivars to yield one high-depth ATCC sample and one high-depth ECACC sample.

### SMAC-seq

For both experimental conditions tested, 1×10^6 MCF-7 cells (ATCC) were resuspended in 500 μl of ice-cold Cell Fractionation Buffer (Abcam) and incubated for 10 minutes on ice. Nuclei were pelleted by centrifugation for 3 min at 500g, then resuspended in 200 μl of ice-cold Nuclei Wash Buffer (10 mM Tris pH7.4, 10mM NaCl, 3mM MgCl2, 0.1 mM EDTA). The nuclei were then pelleted by centrifugation for 3 min at 500g and resuspended in EcoGII reaction buffer (1X NEB CutSmart Buffer, 0.3 M sucrose). 200U of EcoGII and 0.6 mM SAM were added and nuclei were incubated either at 37°C for 10 min (condition one) or incubated at 37°C with 1000 rpm agitation for 15 min, with SAM replenished at 7.5 min (condition two). DNA was extracted with the Monarch® Genomic DNA Purification Kit (NEB) according to manufacturer’s instructions with mixing by inversion instead of vortexing. 1 μg of Genomic DNA was prepared for Nanopore sequencing using the Ligation Sequencing Kit (Nanopore LSK110). Samples were sequenced for 72 hr on an r9.4 MinION flowcell on the Nanopore MinION Mk1C.

### Read alignment and variant calling

Nanopore reads were aligned to hg38 via minimap2 2.17[23] with parameters -a -x map-ont --cs:long --MD. Illumina reads were mapped to hg38 via bwa mem2 2.0pre2 (sse4.1) with default parameters and duplicate reads were marked via the MarkDuplicates tool in Picard 2.23.8(http://broadinstitute.github.io/picard).

### Phasing

To inform phasing, variants were detected in the MCF-7 Illumina data using HaplotypeCaller in GATK 4.1.9.0[24]. The resulting VCF was phased using whatshap 1.0[18] and aligned nanopore reads. Nanopore reads were then tagged with haplotypes using the ‘haplotag’ function of whatshap.

### Methylation calling

Basecalling along with modified base calls was done using megalodon 2.2.9 with guppy 4.4.0 using the res_dna_r941_prom_modbases_5mC_v001 model for 5mCG detection (PromethION), or the res_dna_r941_min_modbases-all-context_v001 model for 6mA detection (SMAC/MinION).

### Implementation

Methylartist is implemented in Python using SQLite[25], matplotlib[26], seaborn[27], numpy[28], scipy[29], pandas[30], scikit-bio[31], pysam[32](https://github.com/pysam-developers/pysam), bx-python(https://github.com/bxlab/bx-python), and the ONT fast5 API(https://github.com/nanoporetech/ont_fast5_api). Methylartist is available at https://github.com/adamewing/methylartist or via *pip install methylartist*

Command-line arguments to methylartist for all figures presented in this manuscript are available in supplemental materials. Additional documentation and examples are available at https://github.com/adamewing/methylartist.

## Supporting information

Supplemental Information: Command-line arguments

## Ethics approval and consent to participate

Not applicable.

## Data Availability

The dataset supporting the conclusions of this article is available in the NCBI Short Read Archive (SRA) repository as BioProject PRJNA748257.

## Acknowledgements

The authors thank members of the Mater Research Genome Plasticity and Disease group, in particular G. Faulkner and P. Gerdes for testing methylartist and P. Carreira for technical assistance, as well as the Kinghorn Centre for Clinical Genomics for providing PromethION sequencing services and Macrogen Oceania for providing Illumina sequencing services. The authors acknowledge the Translational Research Institute (TRI) for research space, equipment, and core facilities that enabled this research.

## Contributions

SWC cultured cells, carried out SMAC-seq, and tested methylartist. MK cultured cells and extracted DNA for PromethION sequencing. ADE wrote methylartist and wrote the manuscript with input and contributions from all authors.

## Supplemental Figures

**Supplemental Figure 1.**
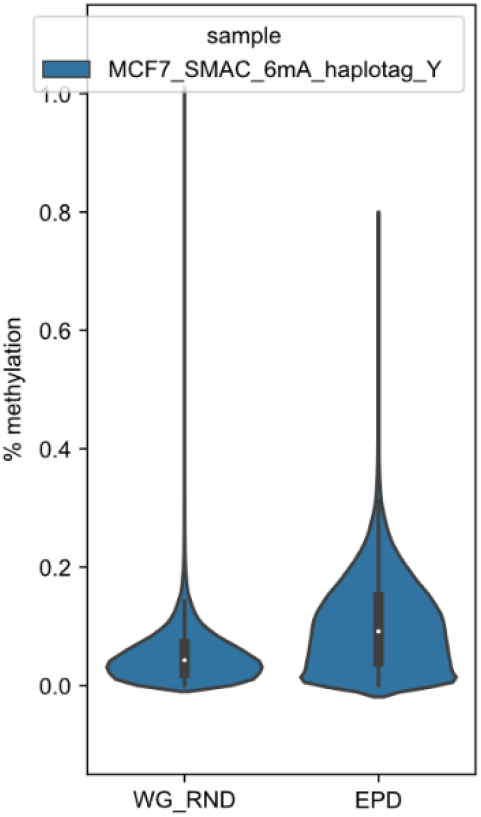
Similar to Figure 1, except for 6mA instead of 5mCG from SMAC-seq on ATCC MCF-7 cells.

**Supplemental Figure 2.**
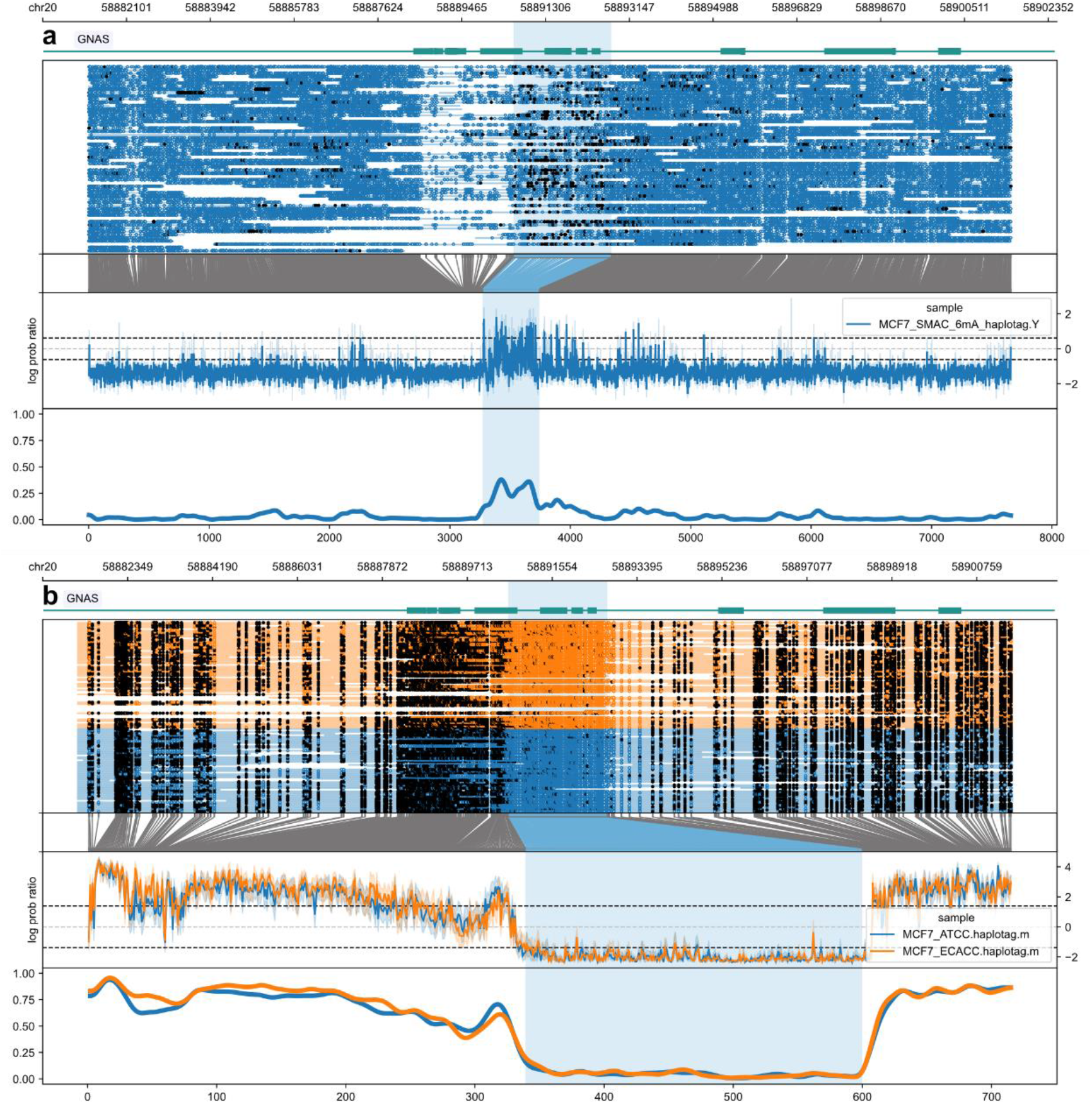
Analogous to Figure 3 but for a regulatory element of the GNAS gene.

## References

1. Zemach A, McDaniel IE, Silva P, Zilberman D. Genome-wide evolutionary analysis of eukaryotic DNA methylation. Science. 2010;328:916–9.

2. Blow MJ, Clark TA, Daum CG, Deutschbauer AM, Fomenkov A, Fries R, et al The Epigenomic Landscape of Prokaryotes. PLoS Genet. 2016;12:e1005854.

3. Couturier M, Lindås A-C. The DNA Methylome of the Hyperthermoacidophilic Crenarchaeon Sulfolobus acidocaldarius. Front Microbiol. 2018;9:137.

4. van Steensel B, Henikoff S. Identification of in vivo DNA targets of chromatin proteins using tethered dam methyltransferase. Nat Biotechnol. 2000;18:424–8.

5. Shipony Z, Marinov GK, Swaffer MP, Sinnott-Armstrong NA, Skotheim JM, Kundaje A, et al Long-range single-molecule mapping of chromatin accessibility in eukaryotes. Nat Methods. 2020;17:319–27.

6. Lee I, Razaghi R, Gilpatrick T, Molnar M, Gershman A, Sadowski N, et al Simultaneous profiling of chromatin accessibility and methylation on human cell lines with nanopore sequencing. Nat Methods. 2020;17:1191–9.

7. Simpson JT, Workman RE, Zuzarte PC, David M, Dursi LJ, Timp W. Detecting DNA cytosine methylation using nanopore sequencing. Nat Methods. 2017;14:407–10.

8. Yuen ZW-S, Srivastava A, Daniel R, McNevin D, Jack C, Eyras E. Systematic benchmarking of tools for CpG methylation detection from nanopore sequencing. Nat Commun. Nature Publishing Group; 2021;12:1–12.

9. Oxford Nanopore Technologies. Megalodon. Github; [cited 2021 Jul 12]. Available from: https://github.com/nanoporetech/megalodon

10. Oxford Nanopore Technologies. Oxford Nanopore Technologies Github. [cited 2021 Jul 13]. Available from: https://github.com/nanoporetech

11. De Coster W, Stovner EB, Strazisar M. Methplotlib: analysis of modified nucleotides from nanopore sequencing. Bioinformatics. Oxford Academic; 2020;36:3236–8.

12. Su S, Gouil Q, Blewitt ME, Cook D, Hickey PF, Ritchie ME. NanoMethViz: an R/Bioconductor package for visualizing long-read methylation data. bioRxiv. 2021;2021.01.18.426757.

13. Pryszcz LP, Novoa EM. ModPhred: an integrative toolkit for the analysis and storage of nanopore sequencing DNA and RNA modification data. bioRxiv. 2021;2021.03.26.437220.

14. Begik O, Lucas MC, Pryszcz LP, Ramirez JM, Medina R, Milenkovic I, et al Quantitative profiling of native RNA modifications and their dynamics using nanopore sequencing. bioRxiv. 2021;2020.07.06.189969.

15. Li F, Guo X, Jin P, Chen J, Xiang D, Song J, et al Porpoise: a new approach for accurate prediction of RNA pseudouridine sites. Brief Bioinform. 2021; Available from: http://dx.doi.org/10.1093/bib/bbab245

16. Comşa Ş, Cîmpean AM, Raica M. The Story of MCF-7 Breast Cancer Cell Line: 40 years of Experience in Research. Anticancer Res. 2015;35:3147–54.

17. Huguet EL, McMahon JA, McMahon AP, Bicknell R, Harris AL. Differential expression of human Wnt genes 2, 3, 4, and 7B in human breast cell lines and normal and disease states of human breast tissue. Cancer Res. 1994;54:2615–21.

18. Patterson M, Marschall T, Pisanti N, van Iersel L, Stougie L, Klau GW, et al WhatsHap: Weighted Haplotype Assembly for Future-Generation Sequencing Reads. J Comput Biol. 2015;22:498–509.

19. Murray IA, Morgan RD, Luyten Y, Fomenkov A, Corrêa IR Jr, Dai N, et al The non-specific adenine DNA methyltransferase M.EcoGII. Nucleic Acids Res. 2018;46:840–8.

20. Dreos R, Ambrosini G, Groux R, Cavin Périer R, Bucher P. The eukaryotic promoter database in its 30th year: focus on non-vertebrate organisms. Nucleic Acids Res. 2017;45:D51–5.

21. Ewing AD, Smits N, Sanchez-Luque FJ, Faivre J, Brennan PM, Richardson SR, et al Nanopore Sequencing Enables Comprehensive Transposable Element Epigenomic Profiling. Mol Cell. 2020;80:915–28.e5.

22. Park Y, Wu H. Differential methylation analysis for BS-seq data under general experimental design. Bioinformatics. 2016;32:1446–53.

23. Li H. Minimap2: pairwise alignment for nucleotide sequences. Bioinformatics. 2018; Available from: http://dx.doi.org/10.1093/bioinformatics/bty191

24. DePristo MA, Banks E, Poplin R, Garimella KV, Maguire JR, Hartl C, et al A framework for variation discovery and genotyping using next-generation DNA sequencing data. Nat Genet. 2011;43:491–8.

25. Hipp RD. SQLite. 2020. Available from: https://www.sqlite.org/index.html

26. Hunter JD. Matplotlib: A 2D Graphics Environment. Computing in Science Engineering. 2007;9:90–5.

27. Waskom M. seaborn: statistical data visualization. Journal of open source software. 2021;6:3021.

28. Harris CR, Millman KJ, van der Walt SJ, Gommers R, Virtanen P, Cournapeau D, et al Array programming with NumPy. Nature. 2020;585:357–62.

29. Virtanen P, Gommers R, Oliphant TE, Haberland M, Reddy T, Cournapeau D, et al SciPy 1.0: fundamental algorithms for scientific computing in Python. Nat Methods. 2020;17:261–72.

30. McKinney W. Data Structures for Statistical Computing in Python. Proceedings of the 9th Python in Science Conference. SciPy; 2010. Available from: https://conference.scipy.org/proceedings/scipy2010/mckinney.html

31. The Scikit-Bio Development Team. scikit-bio: A Bioinformatics Library for Data Scientists, Students, and Developers. 2020. Available from: http://scikit-bio.org

32. Li H, Handsaker B, Wysoker A, Fennell T, Ruan J, Homer N, et al The Sequence Alignment/Map format and SAMtools. Bioinformatics. 2009;25:2078–9.

