## Supplemental Information: Command-line arguments for "Methylartist: Tools for Visualising Modified Bases from Nanopore Sequence Data"

**Violin plots (segmeth, segplot)**

5mCG (Figure 1):

```
methylartist segmeth -d MCF7_data_megalodon.txt -i epdNewHuman006.hg38.pad250.rename.hg38.590bp.50krnd.bed -p 32
```

```
methylartist segplot -s epdNewHuman006.hg38.pad250.rename.hg38.590bp.50krnd.MCF7_data_megalodon.segmeth.tsv -v --svg
```

6mA (Supplemental Figure 1):

```
methylartist segmeth -d MCF7_SMAC_data.txt -i epdNewHuman006.hg38.pad250.rename.hg38.590bp.50krnd.bed -p 32 --max_read_density 0.3
```

```
methylartist segplot -s epdNewHuman006.hg38.pad250.rename.hg38.590bp.50krnd.MCF7_SMAC_data.segmeth.tsv -v --svg
```

**Locus Plots (locus)**

Wnt7b (Figure 2a):

```
methylartist locus -d MCF7_data_megalodon.txt -i chr22:45895804-46003016 -g Homo_sapiens.GRCh38.97.chr.sorted.gtf.gz --genes WNT7B --labelgenes --samplepalette viridis --statname log\ prob\ ratio --svg
```

SHH (Figure 2b):

```
methylartist locus -d MCF7_data_megalodon.txt -i chr7:155764089-155848355 -l 155803598-155806098,155807024-155814024 -g Homo_sapiens.GRCh38.97.chr.sorted.gtf.gz --labelgenes --samplepalette viridis --highlightpalette vlag --statname log\ prob\ ratio --svg
```

**Phased Locus Plots (locus --plased)**

TP53INP1 (Figure 2c):

```
methylartist locus -d MCF7_data_megalodon.txt -i chr8:94848291-95050367 -l 94948291-94950367 -g Homo_sapiens.GRCh38.97.chr.sorted.gtf.gz --samplepalette viridis --maskcutoff 1 --labelgenes --phased --genes INTS8,CCNE2,TP53INP1,NDUFAF6 --statname log\ prob\ ratio --svg
```

**Region Plots (region)**

Figure 3:

methylartist region -d MCF7_data_megalodon.txt -p 48 -g Homo_sapiens.GRCh38.97.chr.sorted.gtf.gz -n CG -r Homo_sapiens_assembly38.fasta --skip_align_plot --genepalette viridis --samplepalette viridis -i chr22:1-50818468 --nticks 20 --highlight_bed chr22.highlights --svg

Contents of chr22.highlights (centromere regions):

```
chr22	1	10510000	#444444
chr22	12954788	15054318	#444444
```

**Composite Plots (composite)**

Figure 5:

methylartist composite -b MCF7_ECACC.haplotag.bam -m MCF7_ECACC.megalodon.db --sample MCF7_ECACC.haplotag_m -s L1HS.MCF7_data_megalodon.excl_ambig.segmeth.tsv -f L1HS -r Homo_sapiens_assembly38.fasta -t L1.3.fa -p 32 --blocks L1.3.highlights.bed --plotmean --svg

**SMAC-seq Locus Plots (locus)**

AXIN-2 5mCpG (Figure 4a):

methylartist locus -d MCF7_data_megalodon.txt -i chr17:65550815-65571075 -l 65560000-65562050 -g Homo_sapiens.GRCh38.97.chr.sorted.gtf.gz --samplepalette viridis --genes AXIN2 --labelgenes --statname log\ prob\ ratio --svg

AXIN-2 6mA (Figure 4b):

methylartist locus -d MCF7_SMAC_data.txt -i chr17:65550815-65571075 -l 65560000-65562050 -g Homo_sapiens.GRCh38.97.chr.sorted.gtf.gz --genes AXIN2 --labelgenes --max_read_density 0.3 --maskcutoff 0 --mincalls 4 --statname log\ prob\ ratio --svg

GNAS 5mCpG (Supplemental Figure 2a):

methylartist locus -d MCF7_data_megalodon.txt -i chr20:58881271-58901531 -l 58890600-58892750 -g Homo_sapiens.GRCh38.97.chr.sorted.gtf.gz --genes GNAS --labelgenes --statname log\ prob\ ratio --svg

GNAS 6mA (Supplemental Figure 2b):

methylartist locus -d MCF7_SMAC_data.txt -i chr20:58881271-58901531 -l 58890600-58892750 -g Homo_sapiens.GRCh38.97.chr.sorted.gtf.gz --genes GNAS --labelgenes --max_read_density 0.3 --maskcutoff 0 --mincalls 4 --statname log\ prob\ ratio --svg
